# Altered muscle niche contributes to myogenic deficit in the D2-*mdx* model of severe DMD

**DOI:** 10.1101/2023.03.27.534413

**Authors:** Davi A. G. Mázala, Ravi Hindupur, Young Jae Moon, Fatima Shaikh, Iteoluwakishi H. Gamu, Dhruv Alladi, Georgiana Panci, Michèle Weiss-Gayet, Bénédicte Chazaud, Terence A. Partridge, James S. Novak, Jyoti K. Jaiswal

**Affiliations:** Center for Genetic Medicine Research, Children’s National Research Institute, Children’s National Research and Innovation Campus, Children’s National Hospital, Washington, D.C., 20012, USA; Department of Kinesiology, College of Health Professions, Towson University, Towson, MD, 21252, USA; Department of Biochemistry and Orthopaedic Surgery, Jeonbuk National University Medical School and Hospital, Jeonju, 54907, Republic of Korea; Institut NeuroMyoGène, Unité Physiopathologie et Génétique du Neurone et du Muscle, INSERM U1513, CNRS UMR 5261, Université Claude Bernard Lyon 1, Univ Lyon, Lyon, France; Departments of Pediatrics and Genomics and Precision Medicine, The George Washington University School of Medicine and Health Sciences, Washington, D.C., 20052, USA

**Keywords:** Duchenne muscular dystrophy, D2-mdx, B10-mdx, B6-mdx, myogenic deficit, inflammation, fibrosis, calcification, muscle regeneration, satellite cell, macrophage, macrophage skewing, fibroadipogenic progenitor, FAP

## Abstract

Lack of dystrophin is the genetic basis for the Duchenne muscular dystrophy (DMD). However, disease severity varies between patients, based on specific genetic modifiers. D2-*mdx* is a model for severe DMD that exhibits exacerbated muscle degeneration and failure to regenerate even in the juvenile stage of the disease. We show that poor regeneration of juvenile D2-*mdx* muscles is associated with enhanced inflammatory response to muscle damage that fails to resolve efficiently and supports excessive accumulation of fibroadipogenic progenitors (FAPs). Unexpectedly, the extent of damage and degeneration of juvenile D2-*mdx* muscle is reduced in adults and is associated with the restoration of the inflammatory and FAP responses to muscle injury. These improvements enhance myogenesis in the adult D2-*mdx* muscle, reaching levels comparable to the milder (B10-*mdx*) mouse model of DMD. *Ex vivo* co-culture of healthy satellite cells (SCs) with the juvenile D2-*mdx* FAPs reduced their fusion efficacy and *in vivo* glucocorticoid treatment of juvenile D2 mouse improved muscle regeneration. Our findings indicate that aberrant stromal cell response contributes to poor myogenesis and greater muscle degeneration in dystrophic juvenile D2-*mdx* muscles and reversal of this reduces pathology in adult D2-*mdx* mouse muscle, identifying these as therapeutic targets to treat dystrophic DMD muscles.

## Introduction

Duchenne muscular dystrophy (DMD) is a progressive X-linked myopathy caused by mutations that ablate expression of the muscle structural protein dystrophin that links myofibrillar actin and the extracellular matrix (ECM)^1–4^. Lack of dystrophin makes the myofiber sarcolemma susceptible to injury and compromises sarcolemmal repair, causing asynchronous myofiber damage and chronic inflammation^5–8^. Muscle from DMD patients and animal models display chronic inflammation, ECM remodeling, fibro-fatty replacement, and progressive muscle loss that diminishes muscle function^9–12^. Additionally, there is evidence of progressive reduction of satellite cells (SC) and their myogenic capacity due to constant muscle injury and turnover, which leads to myofiber loss and replacement by fibrotic tissue in DMD patients^13, 14^. Chronic inflammation and ECM degradation also alters the muscle niche that supports SC function, while genetic modifiers that affect the ECM alter disease severity in DMD patients^15–17^.

One of the genetic modifiers of DMD is the polymorphism in latent transforming growth factor binding protein 4 (LTBP4) that diminishes sequestration of transforming growth factor β (TGF-β) in its latent state^18^. Mice of the DBA/2J (D2) background carry a LTBP4 allele that fails to keep TGF-β in its latent state^19^. D2 mice that also lack dystrophin (D2-*mdx*) mimic the more severe disease observed in DMD patients^19–24^. TGF-β modulates dynamic interactions of macrophages, SCs and other muscle interstitial cell types during healthy muscle regeneration^25–29^. Heightened TGF-β activity disrupts the muscle extracellular niche by altering the crosstalk between stromal cells, including inflammatory cells such as macrophages, and fibroadipogenic progenitors (FAPs), whose interactions support regenerative myogenesis^30–32^. Disruption of macrophage and FAP interactions in damaged muscle delays FAP clearance and promotes fibrosis that further impairs myogenesis^29, 30, 33–36^. Chronic inflammation due to recurrent injury disrupts the synchrony of stromal cell communication required for successful muscle repair^6, 8, 33^. The impact of asynchronous muscle reparative response is demonstrated by failed muscle regeneration and fibroadipogenic muscle loss provoked by repeated muscle injury in a milder model of DMD^8, 34^. Spontaneous muscle damage in the severe juvenile D2-*mdx* model is associated with increased degeneration and failed regeneration^22–24^.

Here we demonstrate that the severe muscle damage and myogenic failure observed in juvenile D2-*mdx* is unexpectedly reversed in muscles from adult D2-*mdx*. We performed a comparative analysis of muscle histopathological changes in the adult versus the juvenile D2-*mdx* and examined the underlying mechanism for this difference in disease severity between adult and juvenile D2-*mdx*. This investigation ascribes damage in juvenile muscle to excessive inflammatory and fibroadipogenic response, restoration of which in adult muscle improves regenerative myogenesis. Using *in vivo* injury and *ex vivo* SC-FAP co-cultures, we show that this aberrant stromal interaction drives myogenic deficit in D2-*mdx* muscles and establish the importance of the muscle niche in the regenerative deficit and disease severity in DMD.

## Materials and Methods

### Animals

All animal protocols were reviewed and approved by the Institutional Animal Care and Use Committee (IACUC) of Children’s National Research Institute and Institut NeuroMyoGène. Juvenile (4-7 wk) and adult (8±3 mo) male and female mice were maintained under normal, ambient conditions with continuous access to food/water. Animals were euthanized by CO_2_ with cervical dislocation, and tissues were harvested, frozen and stored at −80°C. For *in vivo* studies, we used dystrophic mouse models harboring a point mutation in Dmd exon 23, C57BL/10ScSn-mdx/J (B10-*mdx*) and DBA/2J-*mdx* (D2-*mdx*) mice, as well as their corresponding genotype controls – C57BL/10ScSnJ (B10-WT) and DBA/2J (D2-WT). For *in vitro* culture studies, primary cells were harvested from C57BL/6-*mdx*/J (B6-*mdx*) and DBA/2J-*mdx* (D2-*mdx*) mice. As SC from DBA2/J mice exhibit intrinsic deficit in their myogenic ability^23^, SC were used from C57BL/6 mice based on our prior finding that C57BL/6 and C57BL/B10 muscles exhibit similar myogenesis^24^. All mice were originally obtained from The Jackson Laboratory and bred in-house for all experiments.

### BrdU Labeling

5′-bromo-2′-deoxy-uridine (BrdU) (Sigma-Aldrich, B9285) was administered ad libitum in drinking water (0.8 mg/mL) and kept protected from light during administration^24, 37^. Mice received BrdU ad libitum in drinking water from 24–72 h after NTX-induced injury. Mice were subsequently euthanized, and tissues were harvested for processing 3 d after cessation of BrdU administration^24^.

### Toxin-induced injury

Animals were anesthetized with isoflurane and the anterior hind limb was shaved before intramuscular injection of notexin (NTX) or cardiotoxin (CTX) as previously described^24, 34^.

### Deflazacort treatment

Deflazacort (1mg/kg, daily, I.P., Sigma-Aldrich, 1166116) was administered to D2-WT mice (4 wk) within 24 h following NTX injury and continued daily for a period of 7 d. Control D2-WT mice were administered saline. Mice received BrdU ad libitum in drinking water from 24–72 h after NTX.

### Histology and immunofluorescence

Frozen muscles were sectioned at 8 μm thickness using a Leica CM1950 cryostat chilled to −20°C, where tissues were mounted on slides and stained using Hematoxylin and Eosin (H&E), Alizarin Red, and Masson’s Trichrome according to TREAT-NMD Standard Operating Procedures (SOPs) for quantification of damage, calcification and fibrosis, respectively, as previously described^24^, or for immunostaining procedures as previously described^24, 37^. Muscle sections were stained with primary and secondary antibodies as described in ***Supplemental Table 1***.

### Microscopy

We used Olympus VS120-S5 Virtual Slide Scanning System with UPlanSApo 40×/0.95 objective, Olympus XM10 monochrome camera, and Olympus VS-ASW FL 2.7 imaging software. Analysis was performed using Olympus CellSens 1.13 and ImageJ software.

### Gene expression

Triceps muscles were used to perform gene expression analysis. Total RNA was extracted from muscle samples by standard TRIzol (Life Technologies) isolation. Purified RNA (400ng) was reverse-transcribed using Random Hexamers and High-Capacity cDNA Reverse Transcription Kit (Thermo Fisher, 4368814). The mRNAs were quantified using individual TaqMan assays described in ***Supplemental Table 2*** on an ABI QuantStudio 7 Real-Time PCR machine (Applied Biosystems) using TaqMan Fast Advanced Master Mix (Thermo Fisher, 4444556).

### TGF-β1 ELISA

Levels of active TGF-β1 in triceps were quantified using Quantikine ELISA mouse TGF-β1 immunoassay (R&D Systems, MB100B) according to the manufacturer’s recommendations and as previously described^24^. Final values were normalized to total protein concentration.

### Isolation of satellite cells and fibroadipogenic progenitors

SCs were isolated from hindlimb muscles of C57BL/6 mice (n=4-6) using negative selection MACS Satellite Cell Isolation Kit (Miltenyi, 130-104-268) according to manufacture’s protocols. Muscles were minced and incubated with Muscle Dissociation Buffer (Ham’s F-10 (Sigma, N6908), 5% horse serum (Gibco, 16050-130), 1% penicillin/streptomycin (P/S) (Gibco, 15140-122), collagenase II (Gibco, 17101-105)) at 37°C for 60 min with agitation (60-70 RPM). Suspensions were re-incubated with collagenase II and dispase (Gibco, 17105-041) in Ham’s F-10 supplemented with 5% Horse Serum and 1% P/S at 37°C with agitation for 30 min. Suspensions were filtered and MACS LS column sorted (Miltenyi, 130-042-401). Cells were re-suspended in DMEM F-12 (Gibco, 31331-028) supplemented with 20% FBS (Gibco, 10270-106), 2% Ultroser G (Pall Gelman Sciences, 15950-017) and 1% P/S. FAPs were isolated from gastrocnemius muscles of n=4-6 juvenile C57BL/6 (4 d post-CTX), juvenile (7-week-old) D2-*mdx*, or adult (22-week-old) D2-*mdx* by pre-plating the cell suspension for 4 h after the digestion procedure described above. Cells were washed with PBS and left to amplify in non-coated flasks for 4-5 days in growth medium (DMEM F-12, 10% FBS, 1% P/S).

### Co-culture of satellite cells and fibroadipogenic progenitors

SCs and FAPs were co-cultured using 0.4 µm porous transwell culture inserts (Nunc, 056408). SCs were plated in the bottom of the transwell coated with HGF Matrigel (BD Biosciences, 354234), while FAPs were plated in the upper insert. SCs and FAPs were plated in 1:3 ratio in low serum growth medium (DMEM, 2.5% FBS, 1% P/S) and assays were performed after 48 h. For proliferation assay, SCs were plated at 2000 cells/cm^2^ and EdU incorporation was performed after 48 h and detected using Click-iT™ EdU Cell Proliferation Kit (Thermo Fisher, C10337). For differentiation assay, SCs were plated at 10,000 cells/cm^2^ and myogenin (Santa Cruz, SC-12732) staining was used to evaluate differentiation. For fusion assay, SC were plated at 50,000 cells/cm^2^ and desmin (Abcam, ab32362) staining was performed to quantify fusion index. RAW data acquired from different experiments was normalized to no FAP controls.

### Statistics

GraphPad Prism 9.2.0 was used for all statistical analyses of data. Statistical analysis was performed using the non-parametric Mann-Whitney test. Data normality was assessed for all statistical comparisons. All p-values less than 0.05 were considered statistically significant; *p < 0.05, **p < 0.01, ***p < 0.001, and ****p < 0.0001. Data plots reported as scatter plot with mean ± SD.

## Results

### Juvenile D2-mdx mice exhibit excessive muscle damage

We have described the sudden onset of histological damage, and rapid disease progression in muscles of juvenile (3-4 wk) D2-*mdx*^24^, and reported triceps as amongst the most severely affected muscles in this model^24^. Consistent with other reports in adult D2-*mdx*^19–22^, we observed extensive networks of endomysial and perimysial fibrosis in triceps of adult (>5 months-old) D2-*mdx* (**Fig. 1A,B**). The interstitial fibrosis nearly doubled between juvenile and adult D2-*mdx* muscles, while only a modest increase was observed between juvenile and adult B10-*mdx* muscles (**Fig. 1A,B**). Despite the large increase in fibrosis in adult D2-*mdx* triceps, macroscopic examination revealed unexpected improvements in pathological features, prompting a detailed histological examination. H&E staining showed reduced spontaneous myofiber damage and infiltrating mononuclear cells in adult D2-*mdx* than in juvenile D2-*mdx*, to levels comparable to the B10-*mdx* muscles (**Fig. 1C,D, Supplemental Fig. 1**). Alizarin red staining identified a notable decrease in areas of myofiber damage and calcified replacement from ∼15% in juvenile D2-*mdx* to <5% in adult D2-*mdx* (**Fig. 1E,F**). Overall, our analysis revealed that while there is progressive increase in endomysial fibrosis from juvenile to adult D2-*mdx*, surprisingly the extent of damage in the adult D2-*mdx* is reduced as compared to the juvenile D2-*mdx* to levels observed in either juvenile or adult B10-*mdx*.

**Fig. 1.**
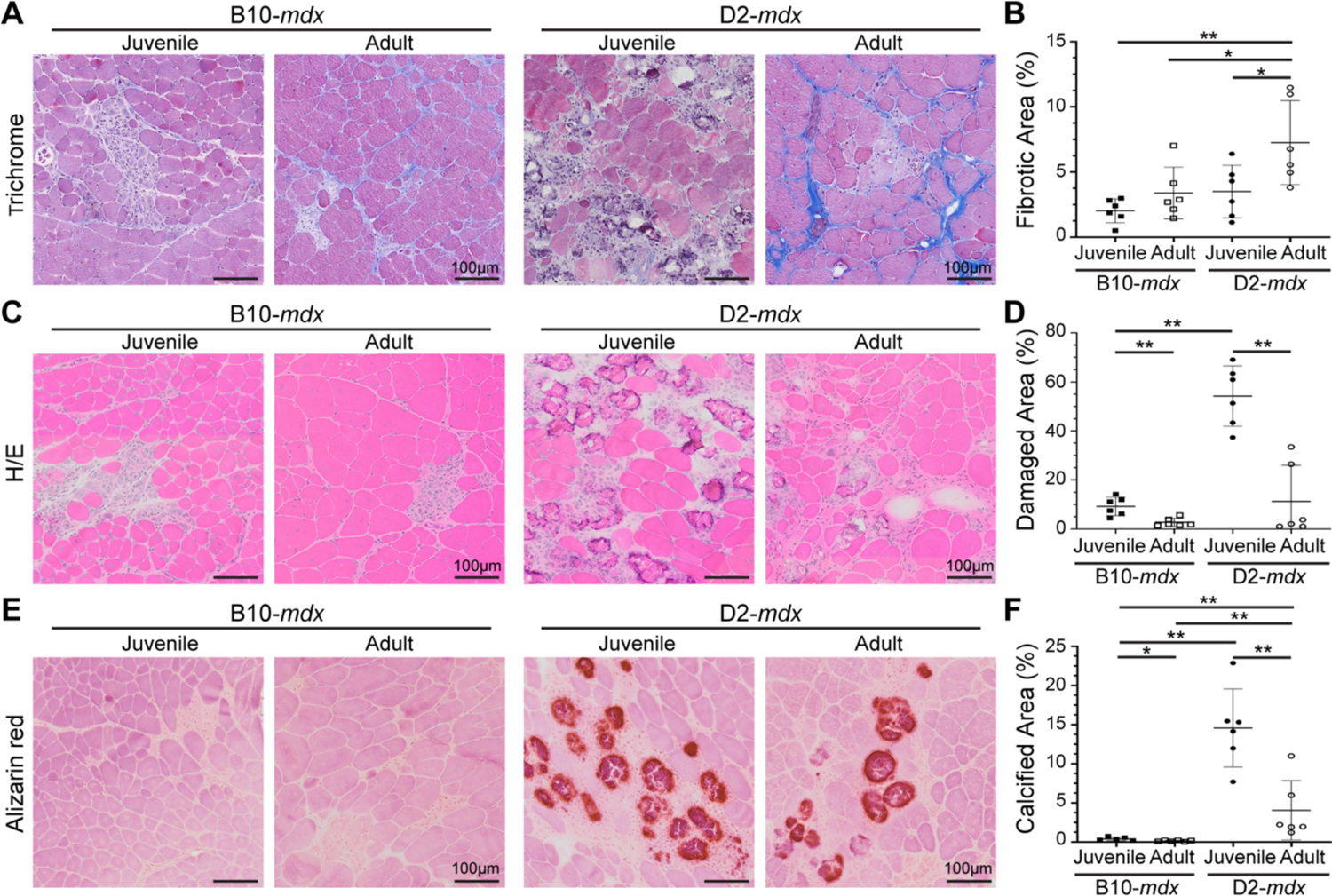
Histopathological assessment of disease in D2-*mdx* and B10-*mdx* models. **A-B.** Masson’s trichrome staining and quantification of percent fibrotic tissue area performed on triceps harvested from juvenile and adult D2-*mdx* and B10-*mdx* mice. **C-D.** H&E staining and quantification of percent damaged muscle tissue area performed on triceps harvested from juvenile and adult D2-*mdx* and B10-*mdx* mice; damaged areas were characterized by the presence of interstitial mononuclear cells, damaged myofibers, and appearance of small-diameter centrally nucleated fibers (CNFs). **E-F.** Alizarin red staining and quantification of percent calcified fiber area performed on triceps harvested from juvenile and adult D2-*mdx* and B10-*mdx* mice. Data represents mean ± SD from n=6 mice per cohort. * *p* < 0.05, ** *p* < 0.01 by Mann-Whitney test. Refer to Supplemental Fig. 1.

### Juvenile D2-mdx muscle exhibits a regenerative deficit that is reversed in adult muscle

When assessing the adult D2-*mdx* histopathology, we observed a notable increase in the frequency of centrally nucleated fibers (CNFs) (**Fig. 1C**). Quantifying myofibers with internal nuclei as a percentage of total myofibers per cross-section revealed nearly 3-times more CNFs in adult D2-*mdx* as compared to juvenile D2-*mdx* (**Fig. 2A,B**). Consequently, while juvenile D2-*mdx* have 5-fold fewer CNFs than juvenile B10-*mdx*, this difference is only 2-fold between the adult D2-*mdx* and B10-*mdx* (**Fig. 2A,B**). As juvenile D2-*mdx* muscles show minimal regenerative ability^21, 23, 24^, we examined if regenerative capacity improved in adult D2-*mdx* muscle, leading to the observed reduction in histopathology (**Fig. 1**). To investigate whether the earlier myogenic deficit in D2-*mdx* is reversed in adulthood, we used notexin (NTX) to acutely injure the tibialis anterior (TA) muscle of D2 wild-type (D2-WT) to avoid confounding effects of chronic muscle injury in D2-*mdx*. To monitor myogenesis that follows this acute *in vivo* injury, we used our 5′-bromo-2′-deoxyuridine (BrdU) ‘myofiber birthdating’ strategy^24, 37^, where BrdU was administered from +1-day to +3-days post injury (dpi) to label regenerated myofibers (**Fig. 2C**). Quantification of total CNFs (**Fig. 2D,E**) and BrdU-labeled CNFs (**Fig. 2D,F**), showed that compared to acutely-injured juvenile D2-WT, regeneration was greatly enhanced in adult D2-WT, mirroring results following spontaneous injury in D2-*mdx* (**Fig. 2A,B**). Nearly 60% of all the myofibers in adult D2-WT muscles were regenerated (CNFs), which was not different from the level of CNFs in B10-WT adult muscle but roughly 3-times greater than in juvenile D2-WT (**Fig. 2D,E**). These numbers mirrored the extent of BrdU-labeled myofibers over 2 dpi, again revealing a similar trend-greater regeneration in adult D2-WT muscles comparable with adult B10-WT. Meanwhile, juvenile D2-WT showed only minor (<5%) BrdU-labeling in randomly dispersed, small-caliber myofibers that constituted large areas of unresolved inflammation even after 6 dpi (**Fig. 2D,F**). Overall, we observed that compared to juvenile D2-WT, there is a greatly improved regenerative response in adult D2-WT muscles (**Fig. 2**).

**Fig. 2.**
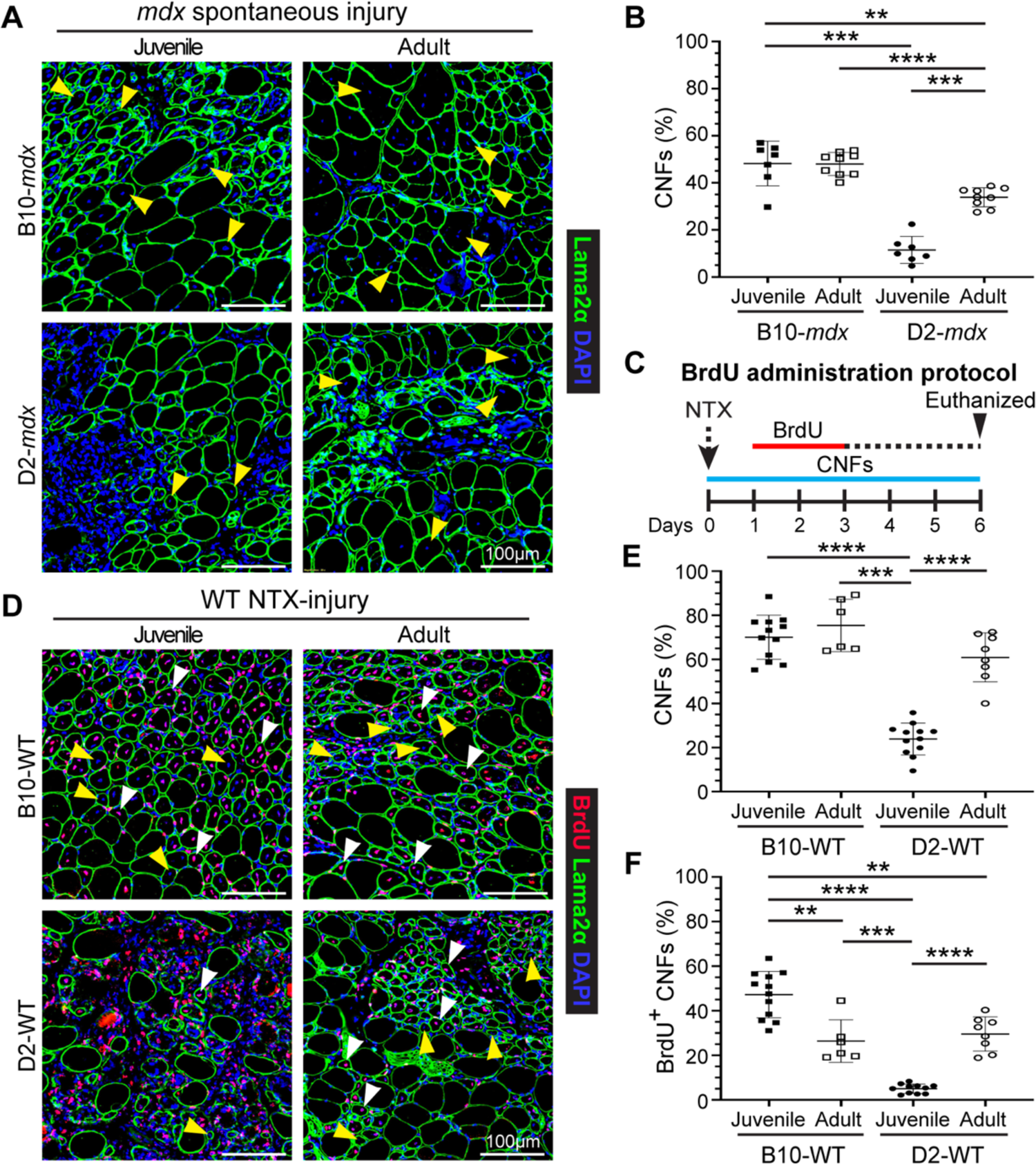
Assessment of muscle regeneration in juvenile and adult *mdx* and WT mice. **A.** IF images from juvenile and adult triceps muscle sections from dystrophic mice stained to identify muscle fibers (Laminin-2α) and CNFs (DAPI). Yellow arrowheads show CNFs. **B.** Quantification of CNFs from dystrophic triceps expressed as a percentage of total muscle fibers. **C.** Schematic showing the BrdU ‘myofiber birthdating’ strategy to label proliferating SCs in NTX-injured TA muscles in juvenile and adult mice by BrdU administration from 24-72h post injury (*green line*). Mice were euthanized and tissues harvested 6d post-injury. **D.** IF images and quantification of muscle sections from B10-WT and D2-WT TA muscles stained to identify muscle fibers BrdU^+^ CNFs and total CNFs; sections co-stained with Laminin-2α and DAPI. White arrowheads show BrdU^+^ CNFs while yellow arrowheads show CNFs. **E.** Quantification of CNFs (%) from NTX-injured TA muscles harvested from juvenile and adult B10-WT and D2-WT mice harvested 6d post-injury. **F.** Quantification of BrdU^+^ CNFs (%) from juvenile and adult B10-WT and D2-WT NTX-injured TA muscles. Data represents mean ± SD from n=7-9 mice per cohort (**B**) or n=6-12 NTX-injured TA muscles per cohort (**E, F**). * *p* < 0.05, ** *p* < 0.01, *** *p* < 0.001, **** *p* < 0.0001 by Mann-Whitney test.

### Stromal alterations mark the regenerative deficit of juvenile D2-mdx muscles

Skeletal muscle regeneration is a multicellular response where SCs interact with the ECM, macrophages, and FAPs to regulate SC proliferation, differentiation, and fusion. To assess the involvement of SC, macrophage, or FAP dysregulation in the regenerative deficit observed in the juvenile D2-*mdx* muscles, we examined the expression of genes associated with these different cell types in triceps (**Fig. 3**). Analysis of the activated SC marker - myoblast determination protein 1 (MyoD), showed that the robust myogenesis observed in B10-*mdx* muscle, was associated with higher levels of MyoD transcript, while the increased damage and regeneration in juvenile (as compared to adult) *mdx* mouse muscle, was associated with higher myogenin (MyoG) transcript in these muscles (**Fig. 3A,B**). To assess whether improvements in regeneration in adult D2-*mdx* was a consequence of increased SCs, we monitored total levels of Paired Box 7 (Pax7) transcript; however, we observed no consistent strain- or age-specific difference for Pax7 transcript (**Fig. 3C**).

**Fig. 3.**
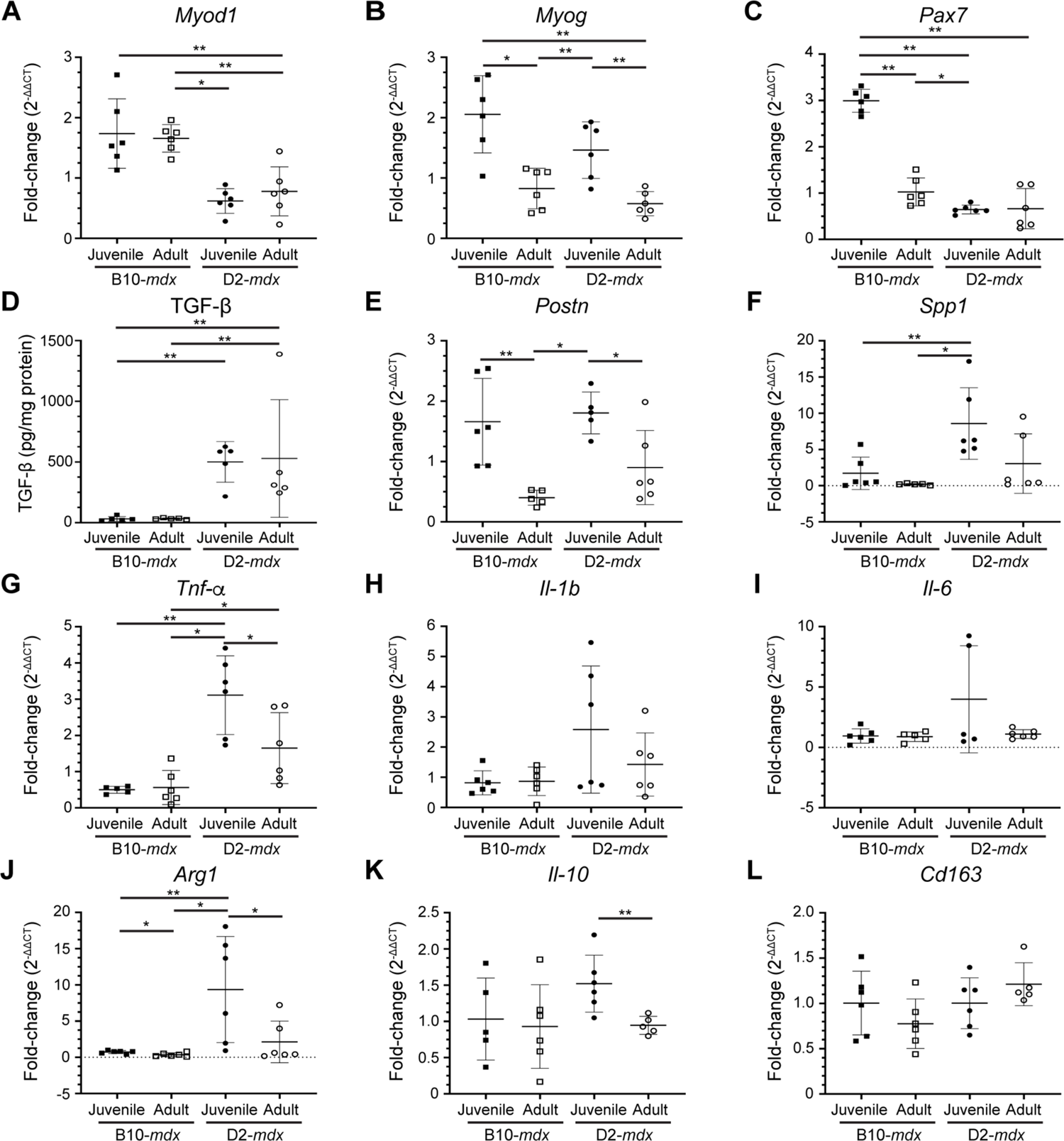
Expression of genes indicative of myogenesis, ECM, and inflammation in juvenile and adult mouse muscles. **A-C.** Gene expression analysis of myogenic markers, MyoD, MyoG and Pax7 in juvenile and adult D2-*mdx* and B10-*mdx* triceps. **D.** Levels of active TGF-β protein assessed by ELISA from juvenile and adult D2-*mdx* and B10-*mdx* triceps. **E-F.** Gene expression analysis of ECM remodeling markers, Postn and Spp1, in juvenile and adult D2-*mdx* and B10-*mdx* triceps. **G-L.** Gene expression analysis of inflammatory genes associated with pro-inflammatory (*Tnf-α, Il-1b, Il-6*) and pro-regenerative (*Arg1*, *Il-10, Cd163*) macrophage phenotypes in juvenile and adult D2-*mdx* and B10-*mdx* triceps. Data represents mean ± SD from n=5-6 mice per cohort. * *p* < 0.05, ** *p* < 0.01 by Mann-Whitney test.

We thus turned to examine FAP/ECM related markers and their dynamics with age and disease progression. TGF-β serves as a master modulator of ECM remodeling and composition during muscle repair and we previously demonstrated its heightened activity in juvenile D2-*mdx* at disease onset^24^. We observed higher TGF-β protein activity in the D2-*mdx* as compared to B10-*mdx*, however, the TGF-β activity levels did not change between juvenile and adult D2-*mdx* (**Fig. 3D**). Due to the extensive effects TGF-β exerts on the regulatory and structural components of the ECM, we next assessed the expression of TGF-β responsive matrix components implicated in dystrophic muscle pathogenesis. Periostin (*Postn*) is a fibroblast-secreted ECM regulatory and structural component whose activity is linked with fibrosis and myogenic function in dystrophic muscle^38^. Like MyoG, greater muscle damage seen in juvenile mice was associated with greater levels of periostin (*Postn*), and this was the same in both D2-*mdx* and B10-*mdx* (**Fig. 3E**). Osteopontin (*Spp1*), a known genetic modifier in DMD patients, functions to influence ECM architecture and fibrosis, while *Spp1* ablation improves muscle function and influences ECM and macrophage polarization^39–43^. In contrast to *Postn*, *Spp1* was upregulated in juvenile D2-*mdx* compared to adult D2-*mdx,* but this was not the case between B10-*mdx* cohorts (**Fig. 3F**). This suggests that low expression of *Spp1* in adult D2-*mdx* may improve regenerative capacity by regulating macrophage polarization^40, 41^.

Examination of markers of macrophage activity and polarization in B10-*mdx* and D2-*mdx* muscles identified a consistent upregulation of both pro-inflammatory and pro-regenerative macrophage markers in juvenile D2-*mdx* muscles. In terms of pro-inflammatory macrophage markers, while only Tumor necrosis factor alpha (*Tnf-α*) was significantly altered between juvenile and adult D2-*mdx*, both Interleukin 1b (*Il-1b*) and Interleukin 6 (*Il-6*) *e*xhibited elevated expression in juvenile D2-*mdx*, which were restored to B10-*mdx* levels in the adult D2-*mdx* muscles (**Fig. 3G-I)**. Similarly, markers of pro-regenerative macrophages, specifically Arginase 1 (*Arg1*), and Interleukin-10 (*Il-10*), but not Cluster of differentiation 163 (*Cd163*), were significantly increased in juvenile D2-*mdx* compared to adult D2-*mdx* (**Fig. 3J-L**).

Thus, while we observed no consistent change in SC and ECM markers between the juvenile and adult D2-*mdx* or between the juvenile and adult B10-*mdx*, we observe consistent dysregulation of *Spp1*, and inflammatory markers corresponding to both pro-inflammatory and pro-regenerative macrophages in juvenile D2-*mdx* muscle as compared to B10-*mdx* and adult D2-*mdx* (**Fig. 3F-I**). This implicates changes in the muscle inflammatory niche in the poor myogenic response specific to juvenile D2-*mdx* muscles. Analysis of the local muscle niche requires spatial exploration of the inflammatory response to monitor the histologically defined damaged regions of the muscle.

### Regenerative deficit of juvenile D2-mdx is linked to heightened pro-inflammatory response

The dynamic interplay between pro-inflammatory and pro-regenerative macrophages is critical for timely resolution and repair of the muscle tissue. To examine the inflammatory response to spontaneous injury of *mdx* muscle we used the pan macrophage marker, F4/80, in conjunction with pro-inflammatory (iNOS) and pro-regenerative (CD206) macrophage markers, to quantify the proportions of pro-inflammatory and pro-regenerative macrophages at and away from the sites of muscle damage (**Fig. 4**). F4/80 immunostaining shows widespread macrophage infiltration in juvenile D2-*mdx* muscle, which is decreased by more than half in muscles from adult D2-*mdx* (**Fig. 4A,B**). Focusing exclusively on the damaged areas characterized by the presence of interstitial mononuclear cells, damaged myofibers, and appearance of small-diameter CNFs, we observed greater abundance of macrophages resulting in greater density of F4/80 labeled macrophages per unit damaged area in juvenile D2-*mdx* (**Fig. 4C**).

**Fig. 4.**
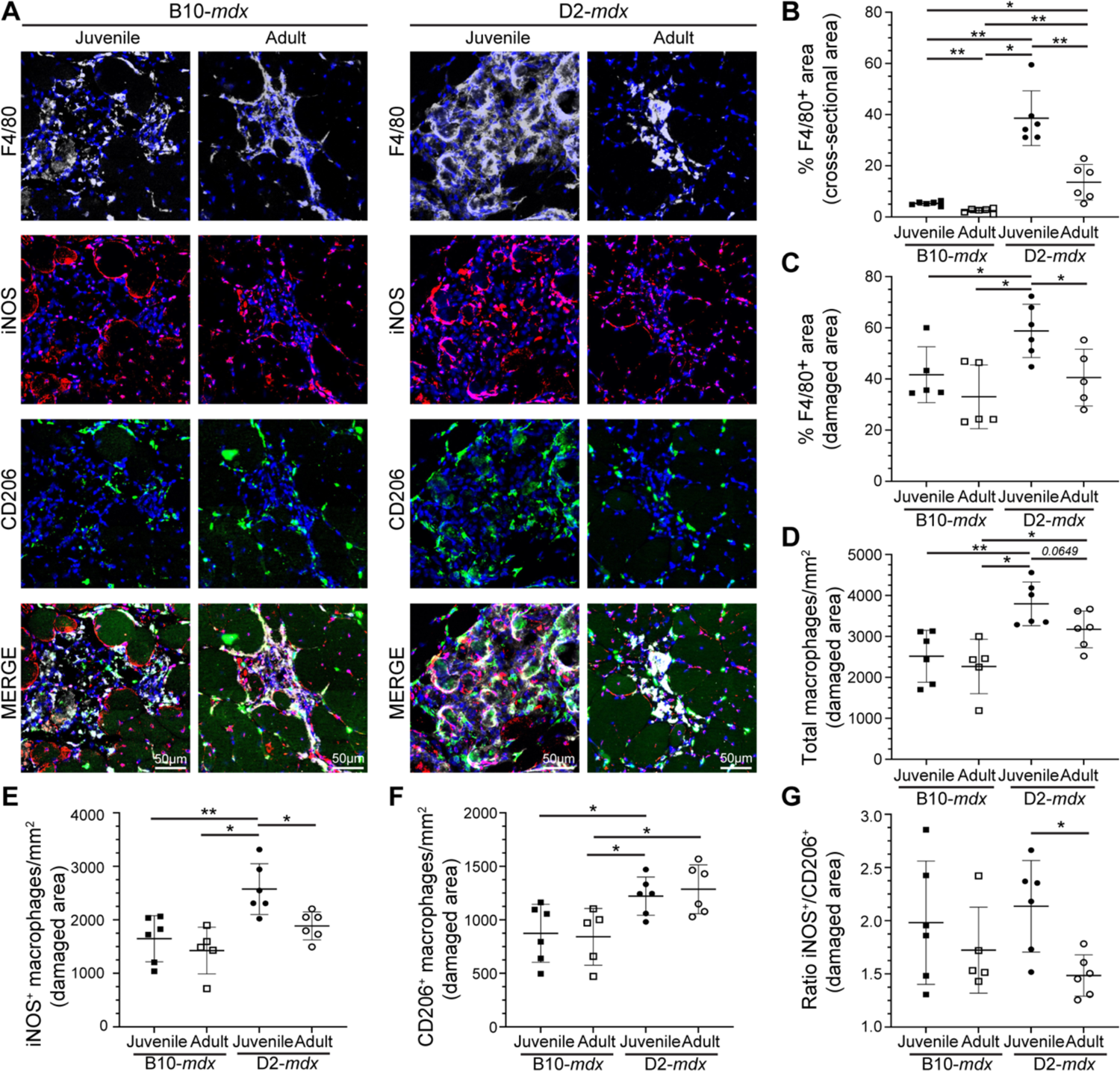
Investigation of macrophage response to spontaneous injury in *mdx* muscles. **A.** Images showing juvenile and adult D2-*mdx* and B10-*mdx* triceps muscle cross-sections stained to mark F4/80, iNOS, and CD206 expressing macrophages. **B-C.** Quantification of F4/80^+^ area (%) per total cross-sectional area (**B**) and only in damaged areas (areas with abundant F4/80^+^ macrophage infiltration) within cross-sections (**C**) in juvenile and adult D2-*mdx* and B10-*mdx* triceps. **D.** Total macrophages per unit area (mm^2^) within damaged regions of juvenile and adult D2-*mdx* and B10-*mdx* triceps cross-sections. **E-G.** Quantification of the distribution of pro-inflammatory (iNOS^+^/F4/80^+^) (**E**), and pro-regenerative (CD206^+^/F4/80^+^) (**F**) macrophages, and the ratio of these macrophages (**G**) within damaged areas of triceps muscles from juvenile and adult D2-*mdx* and B10-*mdx*. Data represents mean ± SD from n=6 mice per cohort. * *p* < 0.05, ** *p* < 0.01 by Mann-Whitney test.

As F4/80 does not distinguish between pro-inflammatory and pro-regenerative macrophages, we next evaluated the contribution of these macrophage subtypes to the total macrophage response observed in juvenile D2-*mdx* by co-labelling tissue sections for iNOS^+^, F4/80^+^ pro-inflammatory, and CD206^+^, F4/80^+^ pro-regenerative macrophages (**Fig. 4A**). The sum of counts of each of these macrophage types in damaged areas corresponded to our finding with (F4/80^+^) macrophage labeling in juvenile D2-*mdx* muscle, where we observed the highest macrophage density per unit damaged area, which was reduced in adult D2-*mdx* muscles (**Fig. 4D**). Monitoring individual macrophage population revealed that the damaged areas of the juvenile D2-*mdx* muscles were enriched in iNOS^+^ and CD206^+^ macrophages. In the adult D2-*mdx* muscle, these pro-inflammatory macrophages in areas of damaged muscle had returned to levels comparable to B10-*mdx,* while the level of pro-regenerative CD206^+^ macrophages remained elevated (**Fig. 4E,F**). Examination of the relative proportion of pro-inflammatory to pro-regenerative macrophages (iNOS^+^/CD206^+^ macrophages), showed that, inflammation in the juvenile D2-*mdx* muscles, relative to adult D2-*mdx* muscle, is skewed towards the pro-inflammatory status (**Fig. 4G**). Together, these analyses indicate that juvenile D2-*mdx* muscles are abnormally inundated with pro-inflammatory macrophages, which correlates with poor myogenic capacity of these muscles.

To assess whether the heightened pro-inflammatory response in juvenile D2-*mdx* is on account of increased entry or greater retention of inflammatory macrophages, we examined the kinetics of the inflammatory response. As *mdx* muscle suffers from spontaneous injuries, and regenerative myogenic deficit is also noted in D2-WT muscle (**Fig. 2E,F**), to achieve a controlled injury scenario we performed acute focal NTX injury to TA muscles of D2-WT and age matched B10-WT and then compared the resulting inflammatory and myogenic response (**Fig. 5-6**). As injury-triggered muscle inflammation progresses from predominantly pro-inflammatory to predominantly pro-regenerative over the week following injury, we monitored total (F4/80^+^) macrophages, as well as levels of pro-inflammatory (iNOS^+^) and pro-regenerative (CD206^+^) macrophages at both an earlier (5 dpi) and later (8 dpi) time point. F4/80 staining at 5 dpi indicated ∼2-fold higher level for juvenile and adult D2-WT than B10-WT counterparts (**Fig. 5A-C**). This indicated that the muscles of D2 mice are predisposed to a stronger inflammatory response irrespective of age. Subsequent assessment of the status of the F4/80 response at 8 dpi showed that the inflammation was largely resolved in the juvenile B10-WT and fully resolved in adult B10-WT and D2-WT muscles (**Fig. 5A-B,D**). In contrast, the extent of inflammation in juvenile D2-WT was much higher than B10-WT, remaining comparable to the levels seen at 5 dpi (**Fig. 5A-C**). Concomitant with the resolution of inflammation in juvenile B10-WT and adult D2-WT muscles at 8 dpi, we observed regenerating myofibers in the site of injury, which were lacking in the juvenile D2-WT muscle, mirroring our earlier observations (**Fig. 2D**). Next, we examined the nature of the macrophages in the areas of inflammation in the acutely injured muscles. Our assessments were limited to 5 dpi as inflammation had resolved by 8 dpi in all cohorts except juvenile D2-WT. We found that both juvenile and adult D2-WT mice mounted a strong inflammatory response that was comparably represented by pro-inflammatory and pro-regenerative macrophages (**Fig. 5E,F**).

**Fig. 5.**
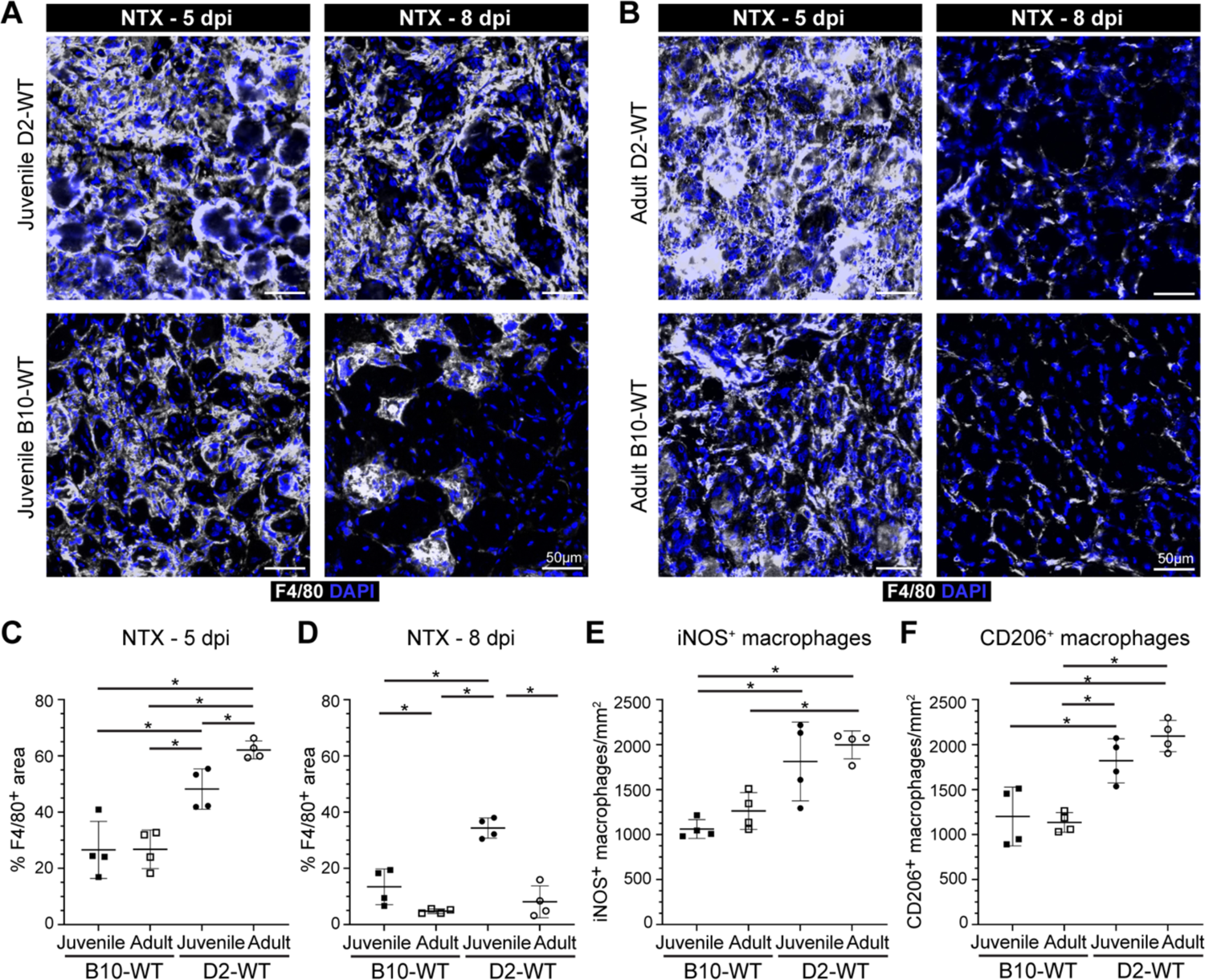
Investigating dynamics of macrophage inflammatory response after acute injury of healthy muscle. **A-B.** F4/80 expression assessed 5 d or 8 d post-injury (dpi) in TA muscles of juvenile (**A**) and adult (**B**) D2-WT and B10-WT mice. IF images of muscle sections stained to show the distribution of macrophages (F4/80) in adult B10-WT and D2-WT NTX-injured TA muscles after 5d and 8d post-injury; muscles co-stained with DAPI. **C-D.** Quantification of F4/80^+^ area after NTX-injury assessed 5 dpi (**C**) and 8 dpi (**D**) in juvenile B10-WT and D2-WT. **E-F.** Quantification of the distribution of pro-inflammatory (iNOS^+^, F4/80^+^) (**E**) and pro-regenerative (CD206^+^, F4/80^+^) (**F**) macrophages per damaged area (mm^2^) in B10-WT and D2-WT NTX-injured TA muscles. Data represents mean ± SD from n=4 mice per cohort. * *p* < 0.05, ** *p* < 0.01 by Mann-Whitney test.

**Fig. 6.**
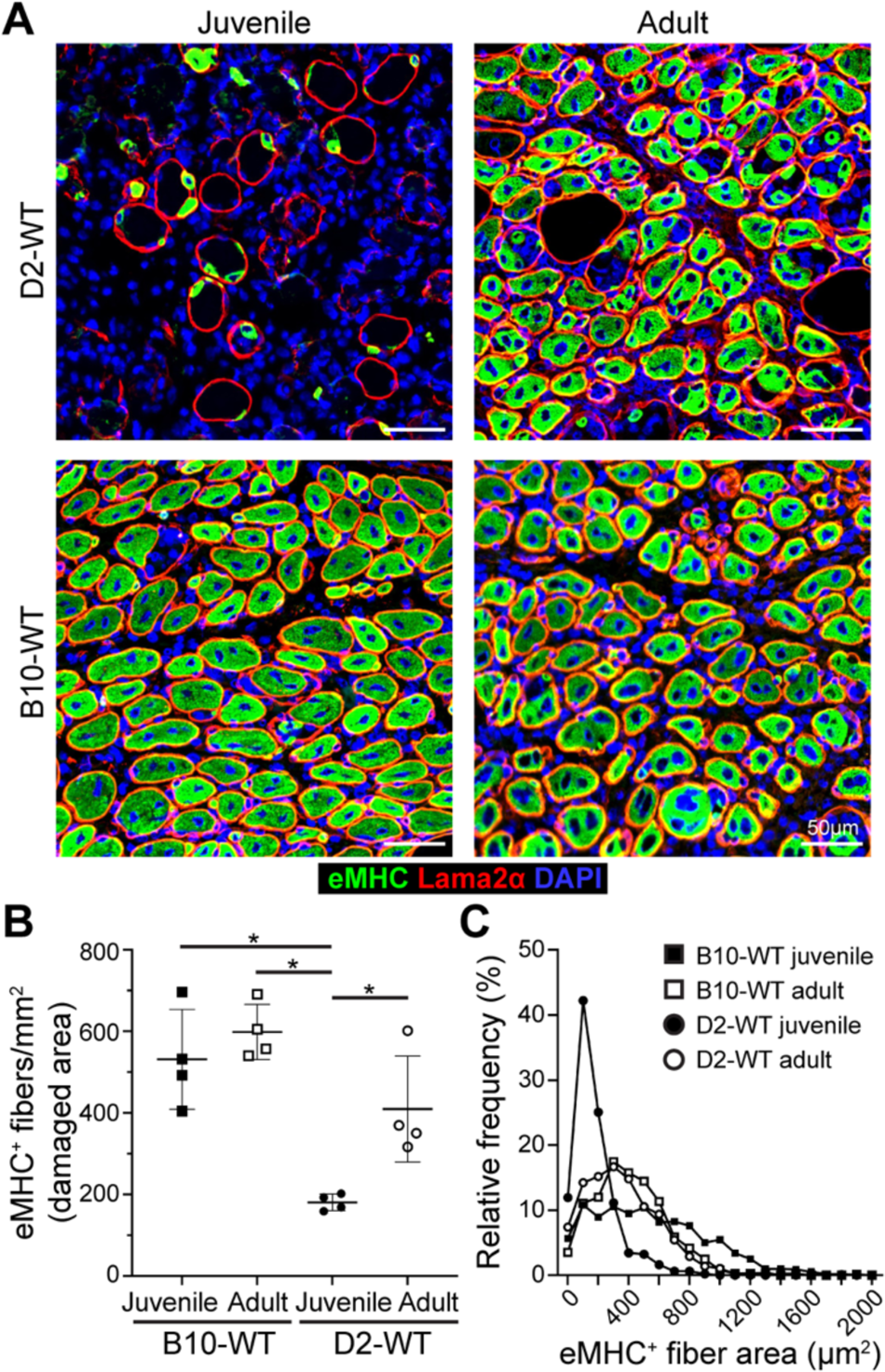
Investigating regenerative response after acute injury of healthy muscle. **A.** eMHC expression assessed 5 d post-injury (dpi) in TA muscles of juvenile and adult D2-WT and B10-WT mice. IF images show the distribution and size of regenerated, eMHC^+^ myofibers (green), co-stained with Laminin-2α (red) and DAPI. **B.** Quantification of eMHC^+^ myofibers per mm^2^ of damaged tissue present 5 dpi in juvenile and adult D2-WT and B10-WT TA muscles. Data represents mean ± SD from n=4 mice per cohort. * *p* < 0.05 by Mann-Whitney test. **C.** Relative frequency plot of eMHC^+^ myofiber area (reported in µm^2^) at 5 dpi in juvenile and adult D2-WT and B10-WT TA muscles, where fiber area was quantified for 1119 fibers (B10-WT juvenile), 496 fibers (D2-WT juvenile), 829 fibers (B10-WT adult) and 866 fibers (D2-WT adult).

As an independent measure for the formation of nascent myofibers we stained acutely injured D2-WT and B10-WT muscles for embryonic myosin heavy chain (eMHC) and monitored these 5dpi in the damaged sites (**Fig. 6**). This showed widespread eMHC expression in small-caliber CNFs throughout the site of injury in all cohorts except juvenile D2-WT (**Fig. 6A, B**). The number of eMHC^+^ fibers in the adult cohort was no different from each other, and the density of eMHC^+^ fibers in adult D2-WT was comparable to juvenile and adult B10-WT muscles 5 dpi, but eMHC^+^ fibers were lacking in juvenile D2-WT muscles (**Fig. 6A, B**). Further, such fibers were notably smaller (< 200 µm^2^) and did not fuse together, even when present within the same basement membrane (**Fig. 6A, C**). Together, these results indicate that D2-WT mice mount a more robust inflammatory response as compared to B10-WT, which fails to resolve in a timely manner in the juvenile D2-WT, leading to the chronic inflammatory response with direct repercussions on regenerative myogenesis.

### FAPs isolated from juvenile D2-mdx mice alter satellite cell fusion capacity in vitro

We previously identified FAP dysregulation is associated with prolonged state of degeneration of D2-*mdx* muscle^24^. Here we examined the role of aberrant stromal response caused by chronic and excessive accumulation of FAPs and inflammatory cells on SC myogenic deficit. We first assessed FAP expansion and numbers during the resolution of spontaneous injury in juvenile D2-*mdx* muscle, by labeling with FAP marker, platelet-derived growth factor receptor-α (PDGFRα). This revealed nearly 2-fold more FAPs in juvenile D2-*mdx* muscle, as compared to the adult D2-*mdx* or the juvenile/adult B10-*mdx* muscle (**Fig. 7A, B**). The observation that FAP abundance in adult D2-*mdx* declines to levels seen in B10-*mdx* muscle suggests that the dysregulated FAP response in the juvenile D2-*mdx* muscles may contribute to the myogenic deficit in these muscles.

**Fig. 7.**
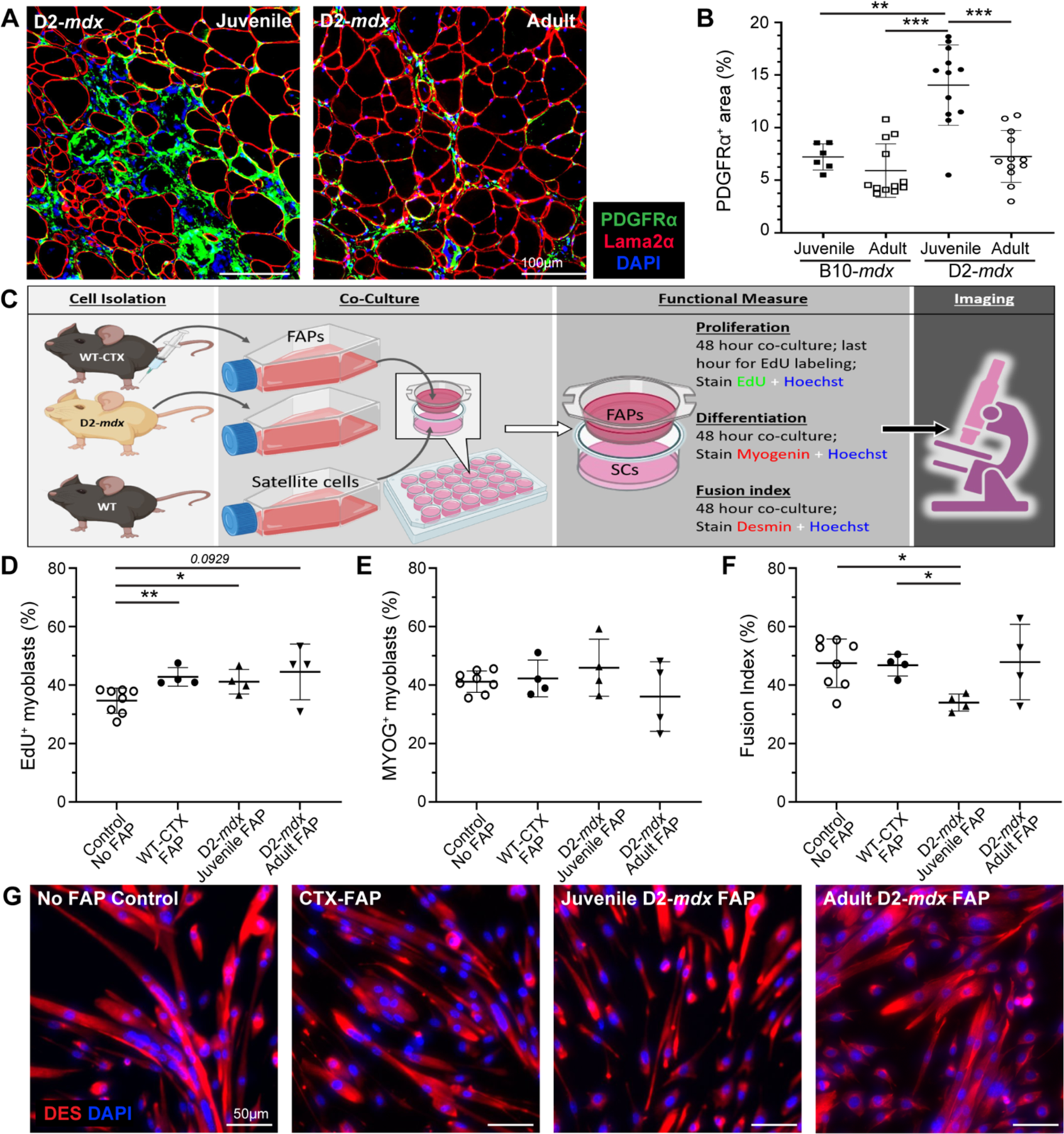
Analysis of juvenile D2-*mdx* FAPs on satellite cell function. **A, B.** IF images and quantification of PDGFRα^+^ FAPs in juvenile and adult D2-*mdx* and B10-*mdx* triceps immunostained for (Laminin), PDGFRα (green), and interstitial nuclei (DAPI). **C.** Schematic illustrating the approach for co-culture experiments to evaluate the effect of FAPs on SC functional properties. SCs from uninjured WT were co-cultured with FAPs isolated from WT mice at 4 days post cardiotoxin injury (WT-CTX), juvenile D2-*mdx* mice, or adult D2-*mdx* mice. **D.** Quantification of the SC proliferation monitored by EdU incorporation. **E.** Quantification of the SC differentiation monitored by Myogenin expression. **F, G.** Quantification of the SC fusion index (**F**) monitored by staining for Desmin expression with corresponding IF images (G) (Desmin stain – red, nuclei stained with Hoechst – blue). Data represents mean ± SD from n=6-12 mice per cohort (A-B) or n=4-6 individual replicates carried out for each functional measure (C-G). * *p* < 0.05, ** *p* < 0.01, *** *p* < 0.001, by Mann-Whitney test.

To investigate whether juvenile D2-*mdx* FAPs impair SC function, we performed co-culture assays and compared the effect of FAPs from juvenile D2-*mdx*, adult D2-*mdx*, and from acutely injured WT mice on the proliferation, differentiation, and fusion of WT SCs (**Fig. 7C**). SCs were plated in the presence of FAPs isolated from either juvenile D2-*mdx* muscles or adult D2-mdx muscles exhibiting spontaneous muscle injury, or from juvenile WT muscles that were acutely injured by cardiotoxin (CTX) (**Fig. 7C-G**). Assessment of proliferation rate of WT SCs by 5’-ethynyl-2’-deoxyuridine (EdU) incorporation showed co-culturing with FAPs enhanced SC proliferation, but no difference in proliferation was observed in co-cultures with the different FAPs - CTX-injured WT, juvenile D2-*mdx*, adult D2-*mdx* (**Fig. 7D**). Next, to examine SC differentiation we quantified the number of myogenin-expressing SCs and found no difference in SC differentiation potential after 48 h when cultured without FAPs or co-cultured with the WT or juvenile or adult D2-*mdx* FAPs (**Fig. 7E**). Finally, we examined fusion capacity of the SCs cultured in the absence of FAPs or in the presence of WT versus D2-*mdx* FAPs harvested from juvenile or adult muscles. This showed a reduction in the fusion index of SCs when co-cultured for 48 h with juvenile D2-*mdx* FAPs, as compared to the no FAP control, CTX-injured WT FAPs, or adult D2-*mdx* FAPs (**Fig. 7F, G**). This final observation recapitulates the above in vivo observation that 5-dpi juvenile D2-WT muscle have the smallest (< 200 µm^2^) nascent myofibers that fail to fuse with the adjacent myofibers. Together, these results identify that poor myogenesis in the juvenile D2 muscles is attributable to a muscle stromal cell niche that inhibits regeneration by inhibiting myotube fusion.

### Glucocorticoid treatment improves myogenesis in D2-mdx muscle

To address whether the altered inflammatory and FAP response specific to juvenile D2-*mdx* muscles is directly responsible for impaired regeneration, we employed an anti-inflammatory glucocorticoid deflazacort treatment regimen in conjunction with acute focal NTX injury to the TA muscles of D2-WT mice to assess potential influence on myogenesis in a controlled injury scenario. Deflazacort (1 mg/kg) treatment was initiated within 24 h of an acute NTX injury and administered daily for 7 d in conjunction with our 3 d (+1 d to +4 d) BrdU-labeling protocol (**Fig. 8A**). Assessment of pro-inflammatory macrophage markers (*Nos2, Il-1b,* and *Il-6*) showed that deflazacort treatment reduced expression of these markers (**Fig. 8B-D**), while pro-regenerative macrophage marker (*Cd163*) was significantly increased relative to controls (**Fig. 8E**). This reflected a change in the macrophage polarization and was associated with reduction in the markers of fibrotic FAPs (*Fn1*, *Col1a1*) in the deflazacort-treated injured muscles (**Fig. 8F-G**).

**Fig. 8.**
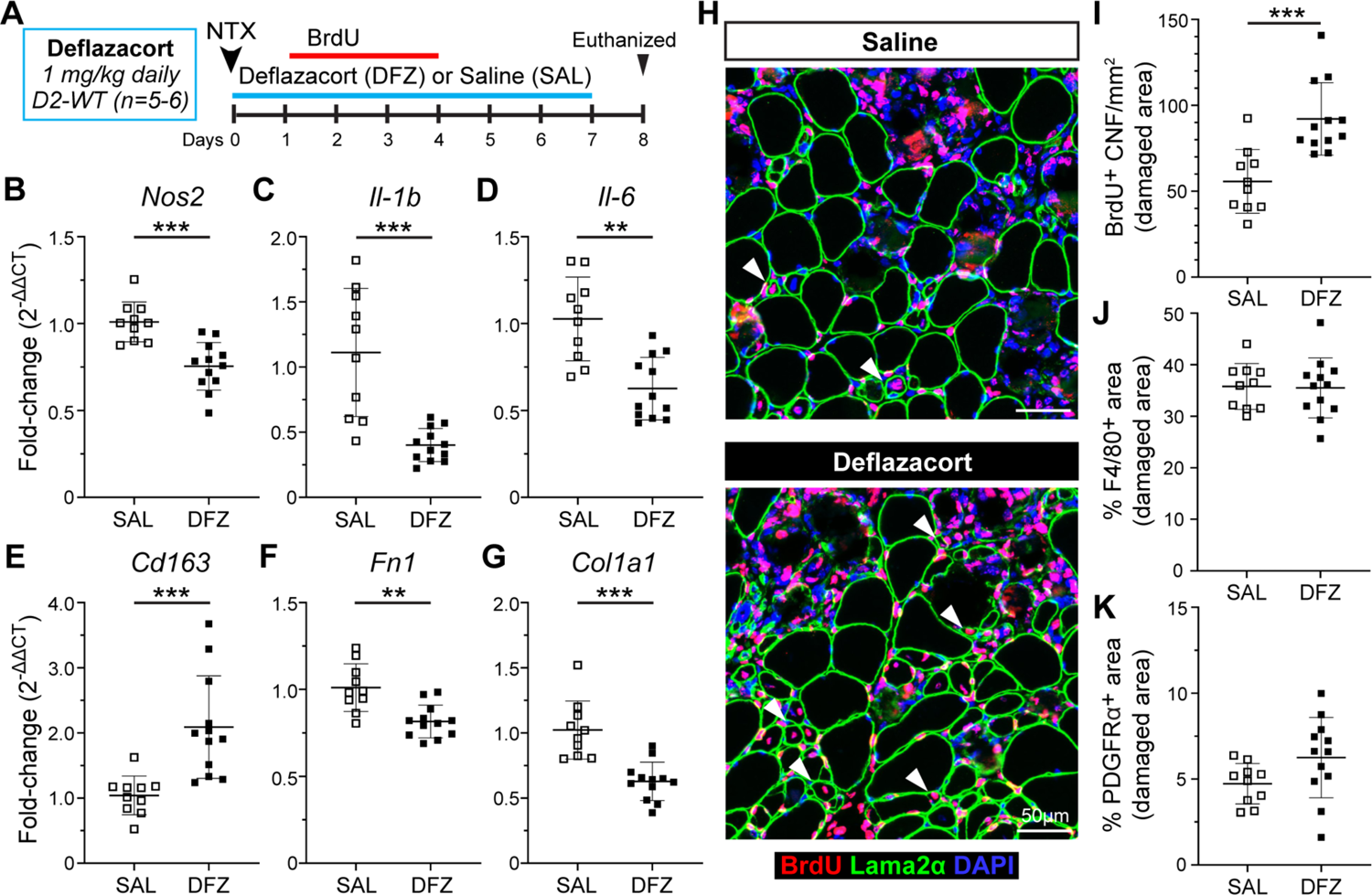
Analysis of regenerative capacity following acute injury and glucocorticoid treatment. **A.** Schematic showing details for deflazacort treatment regimen performed in D2-WT mice following acute NTX injury. For additional details refer to methods. **B-E.** Gene expression analysis of inflammatory genes associated with pro-inflammatory (*Nos2, Il-1b, Il-6*) and pro-regenerative (*CD163*) macrophage phenotypes in juvenile and adult D2-WT and B10-WT TA muscles after acute injury and deflazacort treatment compared to saline controls. **F-G.** Gene expression analysis of ECM markers, *Fn1* and *Col1a1*, in juvenile and adult D2-WT and B10-WT TA muscles after acute injury and deflazacort treatment compared to saline controls. **H.** IF images showing BrdU^+^ CNFs (Red) as indicated by white arrowheads after acute injury and deflazacort treatment compared to saline controls; sections co-stained with Laminin-2α (green) and DAPI (blue). **I.** Quantification of BrdU^+^ CNFs (per damaged area) after acute injury and deflazacort treatment compared to saline controls. **J-K**. % F4/80 (**J**) and PDGFRα (**K**) area reported within damaged muscle regions after acute injury and deflazacort treatment compared to saline controls. Data represents mean ± SD from n=5-6 mice per cohort. ** *p* < 0.01, *** *p* < 0.001 by Mann-Whitney test.

With improvements in macrophage polarization and fibrotic response of the FAPs, we next assessed if these stromal changes caused by glucocorticoid treatment improved regenerative capacity of the juvenile D2 muscles. Analysis of BrdU-labeled CNFs showed that, deflazacort treatment enhanced the myogenic capacity of juvenile D2 muscles, leading to ∼1.6-fold increase in the numbers of regenerated myofibers compared to the control (**Fig. 8H-I**). To evaluate if this improvement in regenerative capacity was the result of reduced macrophage and FAPs within the sites of damage and repair, we also quantified the and PDGFRα^+^ area within the damaged site occupied by macrophages (F4/80^+^) and FAPs (PDGFRα^+^) and observed these were no different between deflazacort-treated and control cohorts (**Fig. 8J-K**). These results indicate the therapeutic potential of glucocorticoid treatment to improve regenerative capacity in juvenile D2 muscles by modulating the macrophage polarization and resulting FAP responses such that the niche created by these stromal cell populations in the injured muscle is more conducive to regenerative myogenesis.

## Discussion

Poor regenerative capacity contributes to DMD severity by limiting the ability of these muscles to effectively replace damaged myofibers lost due to dystrophin deficiency. Like DMD patients, *mdx* mice are characterized by an excessive muscle damage which, in the case of the *mdx* model shows a significant peak during transition from juvenile to adult stage^44^. Here, we aimed to determine the contribution of poor regenerative ability to progressive muscle loss. In the milder *mdx* mouse model, this acute bout of muscle damage is counteracted by robust regenerative myogenesis, which is lacking in the severe D2-*mdx* model^24^. We show that surprisingly, this myogenic deficit in juvenile D2-*mdx* muscles recovers in adult D2-*mdx*, resulting in a greater proportion of centrally nucleated myofibers and greater extent of BrdU incorporation in adult D2-*mdx* muscles than in juvenile D2-*mdx* muscles (**Fig. 2**). Similar to previous studies^22^, we find that the adult D2-*mdx* mice remain less myogenic than the adult B10-*mdx*. However, the improved myogenic ability of the adult D2-*mdx* helps to explain the previous report of amelioration of disease pathology with age in the D2-*mdx* model^20^. It also explains our observation that the extent of muscle damage in adult D2-*mdx* mice is comparable to the less severe B10-*mdx* model (**Fig. 1**).

Disturbances of asymmetric cell division and SC depletion in older individuals have been described as intrinsic impairments in SC that compromise regeneration of the dystrophic muscles^45^. However, we find that the improved myogenesis of the adult D2-*mdx* muscle occurs despite no depletion of SCs in juvenile muscles (indicated by SC-specific markers, Pax7 and MyoD), as compared to adult D2-*mdx* muscle. Concomitantly, expression of myogenin (indicator of myogenic differentiation) is comparable between the mild (B10-*mdx*) and the severe (D2-*mdx*) models (**Fig. 3**). These findings agree with prior work showing comparable SC pool and myogenic activity between dystrophic and WT muscles^46, 47^. Based on *in vivo* SC transplant and *in vitro* analysis of stromal interaction with SCs, it is clear that the muscle niche also plays an important role in SC-mediated regenerative myogenesis^16, 35, 48, 49^. In support of the role played by SC extrinsic factors (muscle niche) in the regulation of myogenesis, we observed that higher expression of ECM and inflammatory regulators including *Spp1*, *Arg1*, *Tnf-α*, *Il-10* is robustly aligned with the regenerative failure observed in the juvenile *D2-mdx* muscle (**Fig. 3**).

Analyses of muscle ECM and inflammatory regulators have established the importance of these factors in regulating SC quiescence, activation and myogenic differentiation^32, 50^. ECM components and stromal cell response to injury has been observed to be altered in the D2-*mdx* mice^19, 24^. In agreement with these changes, we observed a distinct inflammatory response to muscle injury in the D2 (WT and mdx) models, such that juvenile *D2-mdx* muscles exhibit a stronger inflammatory response to injury (**Fig. 4**). In adult *D2-mdx* muscle, the inflammatory response is restored to levels comparable to B10*-mdx*, implicating the excessive inflammatory response in myogenic deficit seen in the juvenile D2-*mdx* mice. Analysis of timed muscle injury in D2-WT mice showed that the excessive inflammatory response to muscle injury in the juvenile D2-WT mice is caused by delayed clearance of inflammatory macrophages that intravasate into the injured tissue (**Fig. 5**), which caused them to adopt an anti-myogenic state hindering regeneration of these inflamed lesions (**Fig. 6**). Such aberrant clearance of inflammatory cells is a hallmark of asynchronous regeneration and was previously implicated in excessive fibrosis and failed regeneration in *mdx* and DMD patient muscles^8, 34^. Concomitant with the co-occurrence of altered ECM and inflammatory responses, we previously demonstrated increased FAP accumulation in the damaged areas of *D2-mdx* muscles, where aberrant FAP responses are inhibitory to regenerative myogenesis^24, 51, 52^. We found that the aberrant stromal (ECM and inflammatory) response alters FAP activity in the juvenile D2-*mdx* such that even in an *ex vivo* co-culture assay, these FAPs significantly suppressed SC-mediated myogenesis. Our analysis determined that it is not the proliferation or differentiation of SCs, but the stage of SC fusion that is diminished selectively by the FAPs derived from the juvenile *D2-mdx* but not from the injured WT muscles or the adult D2-*mdx* muscles (**Fig. 7**). In support of this, we observed that treatment of injured muscles in juvenile D2 mice with deflazacort inhibits the aberrant inflammatory and fibrotic response and improves the stromal cell niche that is more supportive of regenerative myogenesis (**Fig. 8**).

These studies identify the aberrant muscle niche as the driver for myogenic deficit in the juvenile *D2-mdx* model, which is attenuated by maturation of the stromal niche in adult *D2-mdx* muscles, resulting in improved myogenesis in aging animals. This finding suggests targeting the extracellular response to injury as an attractive target to reduce myogenic deficit and severity of disease in DMD.

## Author Contributions

This study was conceived by JSN, TAP, and JKJ. JSN and JKJ designed the experiments with input from DAGM and RH. DAGM, RH, YM and JSN conducted all *in vivo* studies, imaging, and molecular analyses; FS, IHG, and DA assisted RH and JSN in histological assessment and quantification. Cell culture assays were performed by GP, MWG and BC. The manuscript was written by JSN and JKJ, with help from DAGM and RH, and edited by all authors. JSN, BC, and JKJ obtained funding and provided oversight for pursuit of the study.

## Acknowledgements

This work was supported by the National Institutes of Health NIAMS Genetics and Genomics of Muscle Postdoctoral Training Grant (T32AR056993 | DAGM, JSN, JKJ and TAP), the Foundation to Eradicate Duchenne (DAGM, JSN, JKJ), Department of Defense DMDRP (W81XWH2110680 | JKJ; W81XWH2110711| JSN), The Muscular Dystrophy Association (MD480160, MD970000 | JSN), Children’s National Research Institute (CNRI) Institutional Funding (JSN), Towson University College of Health Professions (DAGM), Agence Nationale pour la Recherche (19-CE14-0008-01 | BC) and AFM-Telethon (Alliance MyoNeurALP | BC). Microscopy was performed at the Cell and Tissue Microscopy Core supported by CNRI and The National Institutes of Health NICHD (P50HD105328 | JKJ).

## Availability of Data and Materials

All data will be made promptly available to the scientific community upon request.

---

**Supplemental Table 1.** Primary and secondary antibodies for immunostaining.

| Protein Target | Primary Antibody | Secondary Antibody |
| --- | --- | --- |
| BrdU | Anti-BrdU-biotin, Life Technologies, B35138 (1:100) | Streptavidin Alexa Fluor 488, (1:500), Thermo Fisher, S32354<br>Streptavidin Alexa Fluor 568, (1:500), Thermo Fisher, S11226 |
| eMHC | Anti-eMHC, DSHB, F1.652 (1:25) | Goat anti-mouse IgG1 Alexa Fluor 488, (1:500), Thermo Fisher, A-21121 |
| F4/80 | Anti-F4/80; Bio-Rad, MCA497R (1:100) | Goat anti-rat IgG (H+L) Alexa Fluor 488, (1:500), Thermo Fisher, A-11006<br>Goat anti-rat IgG (H+L) Alexa Fluor 647, (1:500), Thermo Fisher, A-21247 |
| iNOS | Anti-iNOS, Thermo Fisher, PA3-030A (1:100) | Goat anti-rabbit IgG (H+L) Alexa Fluor 488 (1:500), Thermo Fisher, A-11008<br>Goat anti-rabbit IgG (H+L) Alexa Fluor 568 (1:500), Thermo Fisher, A-11011 |
| CD206 | Anti-CD206, Bio-Rad, MCA2235 (1:50) | Mouse Anti-rat IgG2a Alexa Fluor 488 (1:250), Abcam, ab172332 |
| PDGFRα | Anti-PDGFRα, Cell Signaling, D1E1E (1:100) | Goat anti-rabbit IgG (H+L) Alexa Fluor 488 (1:500), Thermo Fisher, A-11008<br>Goat anti-rabbit IgG (H+L) Alexa Fluor 568 (1:500), Thermo Fisher, A-11011 |
| Laminin-α2 | Anti-laminin-α2 (4H8-2); Sigma, L0663 (1:250) | Goat anti-rat IgG (H+L) Alexa Fluor 488 (1:500), Thermo Fisher, A-11006<br>Goat anti-rat IgG (H+L) Alexa Fluor 647 (1:500), Thermo Fisher, A-21247 |

*Description of protein target, primary antibody (manufacturer, catalog number, dilution) and secondary antibody (Alexa Fluor conjugation, manufacturer, catalog number, dilution) shown for all immunostaining procedures performed in this study.*

**Supplemental Table 2.** Taqman assays for quantitative reverse transcriptase PCR (qRT-PCR).

| Gene Target | Gene Name | Taqman Assay |
| --- | --- | --- |
| *Pax7* | Paired box 7 | Mm01354484_m1 [FAM-MGB], Thermo Fisher |
| *Myog* | Myogenin | Mm00446194_m1 [FAM-MGB], Thermo Fisher |
| *Myod1* | Myogenic differentiation 1 | Mm00440387_m1 [FAM-MGB], Thermo Fisher |
| *Spp1* | Secreted phosphoprotein 1 | Mm00436767_m1 [FAM-MGB], Thermo Fisher |
| *Tnf* | Tumor necrosis factor | Mm00443258_m1 [FAM-MGB], Thermo Fisher |
| *Arg1* | Arginase | Mm00475988_m1 [FAM-MGB], Thermo Fisher |
| *Postn* | Periostin | Mm01284919_m1 [FAM-MGB], Thermo Fisher |
| *Il10* | Interleukin 10 | Mm01288386_m1 [FAM-MGB], Thermo Fisher |
| *Nos2* | Nitric oxide synthase 2 | Mm00440502_m1 [FAM-MGB], Thermo Fisher |
| *Il1b* | Interleukin 1 beta | Mm00434228_m1 [FAM-MGB], Thermo Fisher |
| *Il6* | Interleukin 6 | Mm00446190_m1 [FAM-MGB], Thermo Fisher |
| *Cd163* | CD163 antigen | Mm00474091_m1 [FAM-MGB], Thermo Fisher |
| *Fn1* | Fibronectin 1 | Mm01256744_m1 [FAM-MGB], Thermo Fisher |
| *Col1a1* | Collagen type I alpha 1 | Mm00801666_g1 [FAM-MGB], Thermo Fisher |

*Gene symbol, gene name and corresponding Taqman assay for all qRT-PCR procedures performed in this study.*

**Supplemental Fig. 1.**
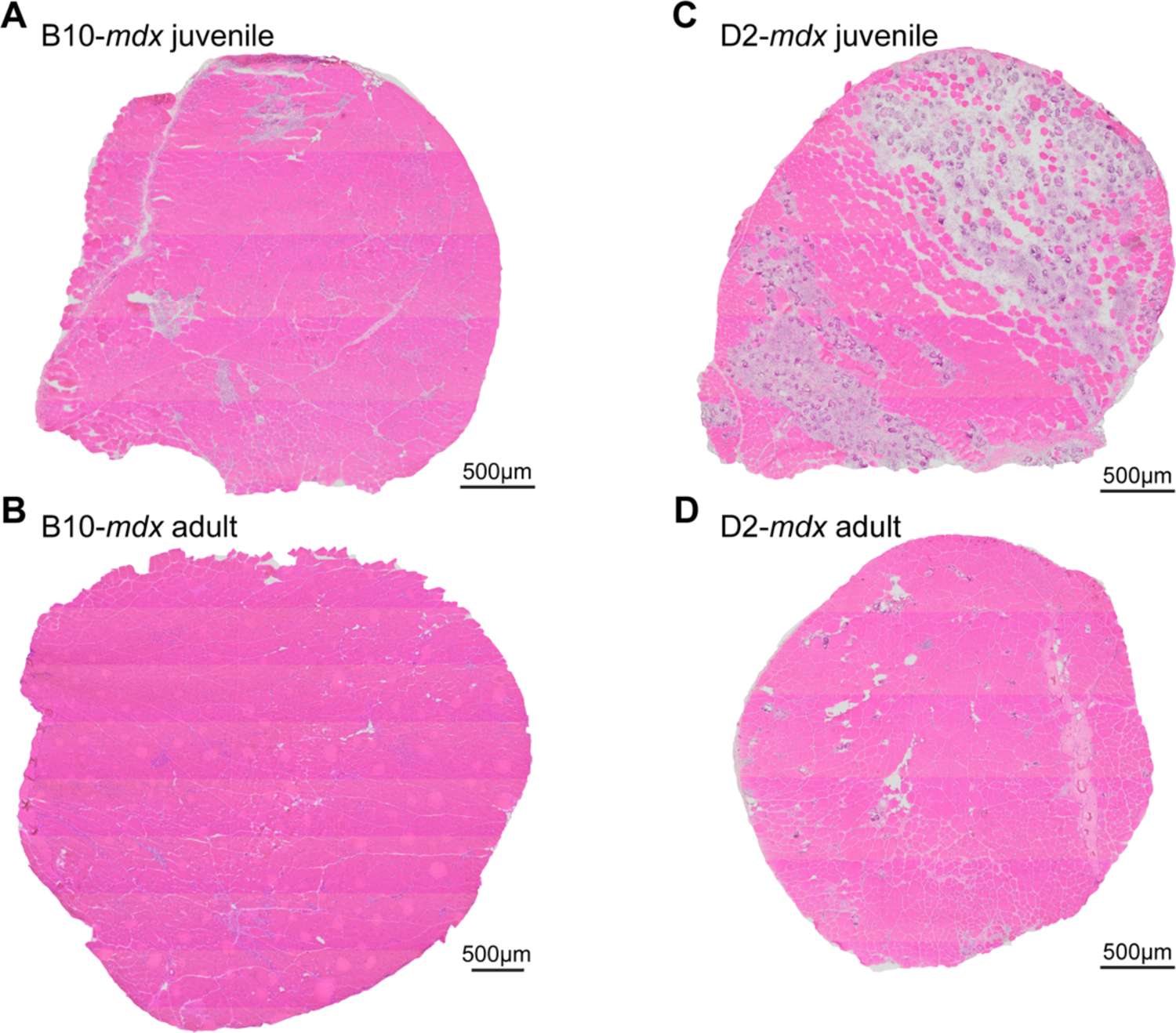
Histopathology in D2-*mdx* and B10-*mdx* models with age. A-D. Whole cross-sectional H&E staining of triceps harvested from juvenile and adult B10-*mdx* (**A**, **B**, respectively) and D2-*mdx* B10-*mdx* (**C**, **D**, respectively) mice.

## References

1. Hoffman EP, Brown RH, Jr., Kunkel LM. Dystrophin: the protein product of the Duchenne muscular dystrophy locus. Cell. 1987;51(6):919–28.

2. Mendell JR, Shilling C, Leslie ND, Flanigan KM, al-Dahhak R, Gastier-Foster J, et al. Evidence-based path to newborn screening for Duchenne muscular dystrophy. Annals of neurology. 2012;71(3):304–13.

3. Ibraghimov-Beskrovnaya O, Ervasti JM, Leveille CJ, Slaughter CA, Sernett SW, Campbell KP. Primary structure of dystrophin-associated glycoproteins linking dystrophin to the extracellular matrix. Nature. 1992;355(6362):696-702.

4. Allikian MJ, McNally EM. Processing and assembly of the dystrophin glycoprotein complex. Traffic. 2007;8(3):177–83.

5. Petrof BJ, Shrager JB, Stedman HH, Kelly AM, Sweeney HL. Dystrophin protects the sarcolemma from stresses developed during muscle contraction. Proceedings of the National Academy of Sciences of the United States of America. 1993;90(8):3710–4.

6. Petrof BJ. The molecular basis of activity-induced muscle injury in Duchenne muscular dystrophy. Mol Cell Biochem. 1998;179(1-2):111–23.

7. Vila MC, Rayavarapu S, Hogarth MW, Van der Meulen JH, Horn A, Defour A, et al. Mitochondria mediate cell membrane repair and contribute to Duchenne muscular dystrophy. Cell Death Differ. 2017;24(2):330–42.

8. Dadgar S, Wang Z, Johnston H, Kesari A, Nagaraju K, Chen YW, et al. Asynchronous remodeling is a driver of failed regeneration in Duchenne muscular dystrophy. The Journal of cell biology. 2014;207(1):139–58.

9. Chen YW, Nagaraju K, Bakay M, McIntyre O, Rawat R, Shi R, et al. Early onset of inflammation and later involvement of TGFbeta in Duchenne muscular dystrophy. Neurology. 2005;65(6):826–34.

10. Kharraz Y, Guerra J, Pessina P, Serrano AL, Munoz-Canoves P. Understanding the process of fibrosis in Duchenne muscular dystrophy. BioMed research international. 2014;2014:965631.

11. Cros D, Harnden P, Pellissier JF, Serratrice G. Muscle hypertrophy in Duchenne muscular dystrophy. A pathological and morphometric study. Journal of neurology. 1989;236(1):43–7.

12. Webster C, Blau HM. Accelerated age-related decline in replicative life-span of Duchenne muscular dystrophy myoblasts: implications for cell and gene therapy. Somatic cell and molecular genetics. 1990;16(6):557–65.

13. Blau HM, Webster C, Pavlath GK. Defective myoblasts identified in Duchenne muscular dystrophy. Proceedings of the National Academy of Sciences of the United States of America. 1983;80(15):4856–60.

14. Decary S, Hamida CB, Mouly V, Barbet JP, Hentati F, Butler-Browne GS. Shorter telomeres in dystrophic muscle consistent with extensive regeneration in young children. Neuromuscular disorders: NMD. 2000;10(2):113–20.

15. Kharraz Y, Guerra J, Mann CJ, Serrano AL, Munoz-Canoves P. Macrophage plasticity and the role of inflammation in skeletal muscle repair. Mediators Inflamm. 2013;2013:491497.

16. Saclier M, Cuvellier S, Magnan M, Mounier R, Chazaud B. Monocyte/macrophage interactions with myogenic precursor cells during skeletal muscle regeneration. The FEBS journal. 2013;280(17):4118–30.

17. Pascual-Morena C, Cavero-Redondo I, Saz-Lara A, Sequi-Dominguez I, Luceron-Lucas-Torres M, Martinez-Vizcaino V. Genetic Modifiers and Phenotype of Duchenne Muscular Dystrophy: A Systematic Review and Meta-Analysis. Pharmaceuticals (Basel). 2021;14(8).

18. Flanigan KM, Ceco E, Lamar KM, Kaminoh Y, Dunn DM, Mendell JR, et al. LTBP4 genotype predicts age of ambulatory loss in Duchenne muscular dystrophy. Annals of neurology. 2013;73(4):481–8.

19. Heydemann A, Ceco E, Lim JE, Hadhazy M, Ryder P, Moran JL, et al. Latent TGF-beta-binding protein 4 modifies muscular dystrophy in mice. J Clin Invest. 2009;119(12):3703–12.

20. van Putten M, Putker K, Overzier M, Adamzek WA, Pasteuning-Vuhman S, Plomp JJ, et al. Natural disease history of the D2-mdx mouse model for Duchenne muscular dystrophy. FASEB journal: official publication of the Federation of American Societies for Experimental Biology. 2019;33(7):8110–24.

21. Coley WD, Bogdanik L, Vila MC, Yu Q, Van Der Meulen JH, Rayavarapu S, et al. Effect of genetic background on the dystrophic phenotype in mdx mice. Human molecular genetics. 2016;25(1):130–45.

22. Hammers DW, Hart CC, Matheny MK, Wright LA, Armellini M, Barton ER, et al. The D2.mdx mouse as a preclinical model of the skeletal muscle pathology associated with Duchenne muscular dystrophy. Scientific reports. 2020;10(1):14070.

23. Fukada S, Morikawa D, Yamamoto Y, Yoshida T, Sumie N, Yamaguchi M, et al. Genetic background affects properties of satellite cells and mdx phenotypes. The American journal of pathology. 2010;176(5):2414–24.

24. Mazala DA, Novak JS, Hogarth MW, Nearing M, Adusumalli P, Tully CB, et al. TGF-beta-driven muscle degeneration and failed regeneration underlie disease onset in a DMD mouse model. JCI Insight. 2020;5(6).

25. Olson EN, Sternberg E, Hu JS, Spizz G, Wilcox C. Regulation of myogenic differentiation by type beta transforming growth factor. The Journal of cell biology. 1986;103(5):1799–805.

26. Cohn RD, van Erp C, Habashi JP, Soleimani AA, Klein EC, Lisi MT, et al. Angiotensin II type 1 receptor blockade attenuates TGF-beta-induced failure of muscle regeneration in multiple myopathic states. Nat Med. 2007;13(2):204–10.

27. MacDonald EM, Cohn RD. TGFbeta signaling: its role in fibrosis formation and myopathies. Curr Opin Rheumatol. 2012;24(6):628–34.

28. Girardi F, Taleb A, Ebrahimi M, Datye A, Gamage DG, Peccate C, et al. TGFbeta signaling curbs cell fusion and muscle regeneration. Nature communications. 2021;12(1):750.

29. Contreras O, Cruz-Soca M, Theret M, Soliman H, Tung LW, Groppa E, et al. Cross-talk between TGF-beta and PDGFRalpha signaling pathways regulates the fate of stromal fibro-adipogenic progenitors. Journal of cell science. 2019;132(19).

30. Lemos DR, Babaeijandaghi F, Low M, Chang CK, Lee ST, Fiore D, et al. Nilotinib reduces muscle fibrosis in chronic muscle injury by promoting TNF-mediated apoptosis of fibro/adipogenic progenitors. Nat Med. 2015;21(7):786–94.

31. Tidball JG. Regulation of muscle growth and regeneration by the immune system. Nat Rev Immunol. 2017;17(3):165–78.

32. Morgan J, Partridge T. Skeletal muscle in health and disease. Dis Model Mech. 2020;13(2).

33. Rosenberg AS, Puig M, Nagaraju K, Hoffman EP, Villalta SA, Rao VA, et al. Immune-mediated pathology in Duchenne muscular dystrophy. Science translational medicine. 2015;7(299):299rv4.

34. Juban G, Saclier M, Yacoub-Youssef H, Kernou A, Arnold L, Boisson C, et al. AMPK Activation Regulates LTBP4-Dependent TGF-beta1 Secretion by Pro-inflammatory Macrophages and Controls Fibrosis in Duchenne Muscular Dystrophy. Cell reports. 2018;25(8):2163–76 e6.

35. Saclier M, Yacoub-Youssef H, Mackey AL, Arnold L, Ardjoune H, Magnan M, et al. Differentially activated macrophages orchestrate myogenic precursor cell fate during human skeletal muscle regeneration. Stem cells. 2013;31(2):384–96.

36. Malecova B, Gatto S, Etxaniz U, Passafaro M, Cortez A, Nicoletti C, et al. Dynamics of cellular states of fibro-adipogenic progenitors during myogenesis and muscular dystrophy. Nature communications. 2018;9(1):3670.

37. Novak JS, Hogarth MW, Boehler JF, Nearing M, Vila MC, Heredia R, et al. Myoblasts and macrophages are required for therapeutic morpholino antisense oligonucleotide delivery to dystrophic muscle. Nature communications. 2017;8(1):941.

38. Lorts A, Schwanekamp JA, Baudino TA, McNally EM, Molkentin JD. Deletion of periostin reduces muscular dystrophy and fibrosis in mice by modulating the transforming growth factor-beta pathway. Proceedings of the National Academy of Sciences of the United States of America. 2012;109(27):10978–83.

39. Vetrone SA, Montecino-Rodriguez E, Kudryashova E, Kramerova I, Hoffman EP, Liu SD, et al. Osteopontin promotes fibrosis in dystrophic mouse muscle by modulating immune cell subsets and intramuscular TGF-beta. J Clin Invest. 2009;119(6):1583–94.

40. Capote J, Kramerova I, Martinez L, Vetrone S, Barton ER, Sweeney HL, et al. Osteopontin ablation ameliorates muscular dystrophy by shifting macrophages to a pro-regenerative phenotype. The Journal of cell biology. 2016;213(2):275–88.

41. Kramerova I, Kumagai-Cresse C, Ermolova N, Mokhonova E, Marinov M, Capote J, et al. Spp1 (osteopontin) promotes TGFbeta processing in fibroblasts of dystrophin-deficient muscles through matrix metalloproteinases. Human molecular genetics. 2019;28(20):3431–42.

42. Hirata A, Masuda S, Tamura T, Kai K, Ojima K, Fukase A, et al. Expression profiling of cytokines and related genes in regenerating skeletal muscle after cardiotoxin injection: a role for osteopontin. The American journal of pathology. 2003;163(1):203–15.

43. Bello L, Pegoraro E. The “Usual Suspects”: Genes for Inflammation, Fibrosis, Regeneration, and Muscle Strength Modify Duchenne Muscular Dystrophy. J Clin Med. 2019;8(5).

44. Coulton GR, Morgan JE, Partridge TA, Sloper JC. The mdx mouse skeletal muscle myopathy: I. A histological, morphometric and biochemical investigation. Neuropathology and applied neurobiology. 1988;14(1):53–70.

45. Chang NC, Chevalier FP, Rudnicki MA. Satellite Cells in Muscular Dystrophy - Lost in Polarity. Trends Mol Med. 2016;22(6):479–96.

46. Boldrin L, Zammit PS, Morgan JE. Satellite cells from dystrophic muscle retain regenerative capacity. Stem Cell Res. 2015;14(1):20–9.

47. Ribeiro AF, Jr., Souza LS, Almeida CF, Ishiba R, Fernandes SA, Guerrieri DA, et al. Muscle satellite cells and impaired late stage regeneration in different murine models for muscular dystrophies. Scientific reports. 2019;9(1):11842.

48. Meng J, Bencze M, Asfahani R, Muntoni F, Morgan JE. The effect of the muscle environment on the regenerative capacity of human skeletal muscle stem cells. Skeletal muscle. 2015;5:11.

49. Joe AW, Yi L, Natarajan A, Le Grand F, So L, Wang J, et al. Muscle injury activates resident fibro/adipogenic progenitors that facilitate myogenesis. Nat Cell Biol. 2010;12(2):153–63.

50. Sousa-Victor P, Garcia-Prat L, Munoz-Canoves P. Control of satellite cell function in muscle regeneration and its disruption in ageing. Nat Rev Mol Cell Biol. 2022;23(3):204–26.

51. Bensalah M, Muraine L, Boulinguiez A, Giordani L, Albert V, Ythier V, et al. A negative feedback loop between fibroadipogenic progenitors and muscle fibres involving endothelin promotes human muscle fibrosis. J Cachexia Sarcopenia Muscle. 2022.

52. Hogarth MW, Uapinyoying P, Mazala DAG, Jaiswal JK. Pathogenic role and therapeutic potential of fibro-adipogenic progenitors in muscle disease. Trends Mol Med. 2022;28(1):8–11.

